# Analyzing the Performance of Deep Learning Splice Prediction Algorithms

**DOI:** 10.1101/2025.11.20.689501

**Authors:** Nathan Fortier, Gabe Rudy, Andreas Scherer

## Abstract

SpliceAI has become the leading computational tool for predicting splice-altering variants, but restrictive licensing has limited its adoption by commercial clinical laboratories. While open-source reimplementations have emerged with author-reported comparisons, independent benchmarking across diverse datasets is needed to establish their practical equivalence.

We compared the original SpliceAI algorithm against two open-source alternatives (OpenSpliceAI and CI-SpliceAI) across three independent benchmarks: a curated dataset of 1,316 functionally validated variants, 213 variants with experimental splice assay data, and 58,064 clinically classified variants from ClinVar. All deep-learning methods were also compared with a legacy ensemble of four traditional splice prediction algorithms (MaxEntScan, NNSplice, GeneSplicer, and PWM), enabling direct comparison between modern and conventional approaches. Across all benchmarks, the deep-learning models consistently outperformed the ensemble of traditional algorithms. We evaluated sensitivity, specificity, and balanced accuracy for each algorithm, and performed statistical testing to assess significance of performance differences. Additionally, we conducted a correlation analysis on 100,000 variants to quantify the concordance of splice-scores and the agreement on splice site positions between implementations.

All three deep learning algorithms demonstrated comparable performance on the literature-curated benchmark (balanced accuracies: 89.5-90.7%) and the ClinVar dataset (88.9-89.5%). While both open-source solutions achieved a statistically significantly higher accuracy than SpliceAI on the ClinVar dataset, the magnitude of this improvement was small and unlikely to be of practical significance. On the functional splice assay dataset, the original SpliceAI achieved the highest accuracy (83.6%), while OpenSpliceAI showed significantly lower performance (74.6%, p = 0.019). Correlation analysis revealed that CI-SpliceAI maintained balanced concordance across splice event types (ρ = 0.786-0.883), whereas OpenSpliceAI exhibited asymmetric performance with stronger correlation for loss events (ρ = 0.924-0.940) than gain events (ρ = 0.668-0.677). Both implementations demonstrated high spatial agreement with SpliceAI, with exact splice site position match rates exceeding 90% for all event types. Together, these results demonstrate that both open-source reimplementations of SpliceAI successfully reproduce the predictive behavior of the original algorithm across multiple evaluation contexts, while consistently outperforming traditional splice prediction methods.

## Introduction

Aberrant splicing is a major contributor to human disease, with an estimated 15–30% of all disease-causing variants having the potential to disrupt normal splicing [1]. While splice-altering variants that disrupt the canonical AG and GT splice dinucleotides are easily identified, it is common for non-coding and missense variants lying outside of these dinucleotides to disrupt normal splicing. The determination of whether a given variant will disrupt an existing splice site or introduce a novel splice site, has proven to be a computationally difficult problem.

Early splice prediction methods attempted to model splice motifs probabilistically. SpliceSiteFinder, one of the first widely adopted tools, used a simple position weight matrix to characterize the nucleotide probabilities surrounding splice sites [2]. GeneSplicer improved upon this by applying markov models and maximal dependence decomposition to better capture dependencies within the motif [3]. Another commonly used method, MaxEntScan, employed maximum entropy modeling to approximate the distributions underlying true splice sites [4].

Neural network–based approaches soon followed. NNSplice represented the first published attempt to use machine learning for this task, applying a shallow feed-forward neural network with a single hidden layer [5]. The introduction of SpliceAI marked a significant advancement in splice site prediction, leveraging deep convolutional neural networks and large genomic contexts. Using a 10,000 base pair window around each candidate splice site, SpliceAI demonstrated substantially improved accuracy over earlier probabilistic and machine-learning models [6].

Since its publication, SpliceAI has become the leading tool for splice site prediction. However, the restrictive licensing surrounding its pre-trained weights and large precomputed score datasets has hindered broader integration into pipelines used for clinical testing. In response, several authors have developed open-source reimplementations and updates to the original model.

OpenSpliceAI is one such open-source method. This approach is a complete reimplementation and retraining of SpliceAI using the PyTorch machine learning library [7]. Another open-source alternative to SpliceAI is CI-SpliceAI, which offers a reimplementation of the algorithm based on an updated version of the TensorFlow library and a retrained model using a collapsed isoform set representative of all manually annotated constitutive and alternative splice sites from GENCODE [8].

In this study, we perform a comprehensive comparison of SpliceAI, OpenSpliceAI, CI-SpliceAI, and a voting ensemble of four conventional methods using multiple benchmark datasets, including curated functional assays, literature-derived variant sets, and a large-scale dataset derived from clinical variant classifications in ClinVar. Additionally, we assess the quantitative concordance between open-source implementations and the original SpliceAI through detailed correlation analysis of predicted splice scores and positions. Our analysis provides empirical guidance for algorithm selection and establishes whether open-source alternatives can serve as viable replacements for the original SpliceAI in research and clinical applications.

## Methodology

### Splice Prediction Tools

This study compares the performance of four different splice prediction tools:

1. **SpliceAI:** a deep convolutional neural network designed to predict splice junctions directly from primary genomic sequence [6].
2. **OpenSpliceAI:** a full open-source reimplementation of SpliceAI built in the PyTorch deep learning framework [7].
3. **CI-SpliceAI:** an open-source TensorFlow implementation of the SpliceAI algorithm trained using a collapsed isoform representation derived from all manually curated constitutive and alternative splice sites in GENCODE [8].
4. **Legacy Ensemble:** performs an ensemble vote (N-of-4) across the four established splice prediction algorithms provided by the VarSeq genomic analysis software: MaxEntScan, NNSplice, GeneSplicer, and a position weight matrix algorithm based on SpliceSiteFinder [9].

The inclusion of the Legacy Ensemble enables evaluation of how much additional predictive accuracy the deep learning– based SpliceAI models provide relative to conventional splice-site prediction strategies. For all benchmark analyses, the Legacy Ensemble voting threshold providing the highest balanced accuracy was used for classification. This threshold was N > 1 for all datasets, with the exception of the deeply intronic ABCA4 variants in the Riepe Benchmark dataset, for which the optimal threshold was N > 2.

For all deep learning algorithms, the following options were used:

- **Masking** was enabled to disregard gains of established splice sites and losses of non-splice sites.
- **Maximum distance** between the variant and gained/lost splice site was set to 500 bp.

### Performance Benchmarking Methodology

The performance of these algorithms was evaluated across three different datasets. For each dataset, the sensitivity, specificity, and balanced accuracy were computed as follows:

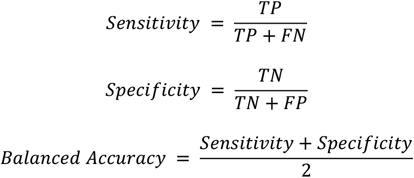

To determine if the observed differences in overall accuracy were statistically significant, a two-proportion Z-test was performed to compare the performance of the open-source models (CI-SpliceAI and OpenSpliceAI) against the original SpliceAI algorithm.

#### CI-SpliceAI Benchmark Dataset

The first dataset comprises 1,316 unique variants aligned to the GRCh37 reference sequence with functionally validated splicing impacts sourced from the literature. This dataset was published by the original CI-SpliceAI authors and is balanced between splice-affecting and non-affecting variants, as well as between strand orientations, reducing potential biases [8].

A binary classification approach was adopted to assess each algorithm’s ability to distinguish between splice-altering and non-splice-altering variants.

- Benchmark variants labeled as alternative acceptor, alternative donor, multiple, retention, or skip were classified as Splice-Altering (Positive).
- Variants labeled normal were classified as Not Splice-Altering (Negative).

For both models, the maximum of the four scores (Acceptor Gain, Acceptor Loss, Donor Gain, Donor Loss) was used. A variant was classified as Splice-Altering (Positive) if the maximum score was > 0.5, corresponding to the recommended threshold for pathogenic classification by SpliceAI [6].

#### Riepe Benchmark Dataset

The second analysis evaluated performance on the functional splice assay dataset published by Riepe *et al*. in 2021, consisting of 152 ABCA4 and 61 MYBPC3 variants [10]. Notably, 81 of the ABCA4 variants are deeply intronic. Functional validation data from LOVD, ClinVar, and ExAC provided precise ground-truth labels distinguishing between specific splice site alterations (acceptor/donor gain or loss).

For this benchmark, the most relevant SpliceAI score was selected based on each variant’s effect type, and score thresholds were optimized per gene and algorithm combination to yield the highest possible accuracy, thus representing an upper-bound estimate of model performance on functional data.

#### ClinVar Benchmark

The final benchmark dataset was constructed from the ClinVar Assessments database (release date: 2025-10-02) [11]. All GRCh38-aligned variants were imported into the VarSeq software and subjected to an initial pre-filtering step in which we removed variants for which none of the four Legacy Ensemble splice prediction algorithms indicated either a gained or disrupted splice site. This step yielded a dataset enriched for variants with at least some computational evidence suggestive of splice relevance.

Following pre-filtering, we applied two independent classification filter chains to assign variants to positive and negative benchmark sets:

- **Splice-Altering (Positive Set):** Variants classified by ClinVar as Pathogenic and whose submitted interpretation or summary text contained the term “splice.” These variants are presumed to have experimentally or clinically supported splice-altering effects.
- **Non–Splice-Altering (Negative Set):** Variants classified by ClinVar as Benign. These variants are assumed not to disrupt normal splicing.

Variants meeting the first criterion were designated as splice-altering, whereas those meeting the second were designated as non-disruptive. This procedure resulted in a combined benchmark dataset of 58,064 variants.

Although this dataset is less stringently curated than the two smaller GRCh37 benchmark datasets, it encompasses a substantially broader spectrum of variant types, enabling direct evaluation of algorithm performance on the GRCh38 reference assembly and across a wide range of genes.

### Correlation Analysis

To quantify the concordance between the scores generated by the open-source algorithms (CI-SpliceAI and OpenSpliceAI) and those from the original SpliceAI algorithm, a correlation analysis was performed. Variants were sampled from the precomputed SpliceAI scores provided by Illumina for the GRCh37 reference genome assembly [6].

Prior to random sampling, the dataset was filtered to exclude variants with minimal evidence of splicing disruption by removing any scores for which all SpliceAI delta scores were below 0.1. A final set of 100,000 variants was randomly sampled from this filtered subset, comprising 50,000 from the masked indel file and 50,000 from the masked SNV (single nucleotide variant) file.

To enable a paired comparison, both the OpenSpliceAI and CI-SpliceAI algorithms were executed to obtain new predictions for the 100,000 variants. Masked delta scores were computed for both algorithms using a maximum distance of 50 bp between the variant and the candidate splice site, thereby matching the parameters used to generate the original precomputed SpliceAI scores.

Four splice disruption categories were evaluated:

- DS_AG: Delta Score Acceptor Gain
- DS_AL: Delta Score Acceptor Loss
- DS_DG: Delta Score Donor Gain
- DS_DL: Delta Score Donor Loss

Each score represents the probability that the variant is splice-altering. The authors recommend a threshold of 0.2 for high recall, 0.5 for balanced accuracy, and 0.8 for high precision. For each score, the following correlation statistics were computed:

- **Spearman’s rank correlation coefficient (ρ):** A non-parametric measure of monotonic association between CI-SpliceAI and SpliceAI scores. Spearman correlation was chosen over Pearson correlation because splice scores are bounded (0-1) and the relative ranking of variants is more clinically relevant than absolute numerical agreement.
- **Mean Absolute Difference:** The average absolute difference between paired scores providing an interpretable measure of typical score divergence.

For variants where either tool predicted a delta score ≥ 0.2 (the high-recall threshold recommended by the SpliceAI authors), we assessed whether the predicted delta positions matched exactly. The delta position (DP) indicates the distance (in base pairs) from the variant to the predicted affected splice site.

For each score type we computed the following:

- **Exact Match Rate:** The percentage of high-confidence predictions for which the delta position matches that reported by SpliceAI.
- **95% Confidence Intervals:** Wilson score intervals for the match rate, accounting for binomial uncertainty.

## Experimental Results

### CI-SpliceAI Benchmark Results

When compared on the CI-SpliceAI benchmark data, all three deep-learning algorithms demonstrated highly similar, strong performance, with overall balanced accuracies exceeding 89%. These results seem to confirm that both algorithms successfully replicate the general predictive capability of the original SpliceAI model.

**Figure 1.**
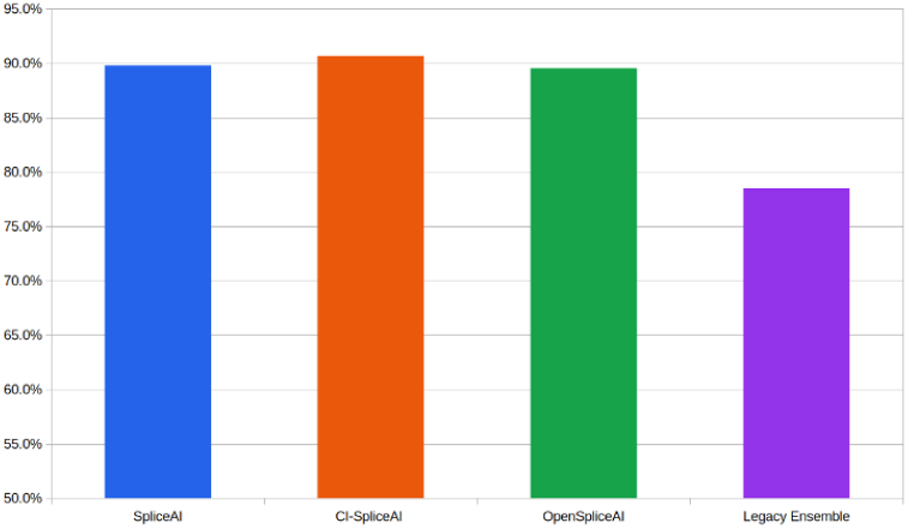
Balanced Accuracy on CI-SpliceAI Benchmark

Although CI-SpliceAI achieved the highest balanced accuracy on this dataset, neither CI-SpliceAI (two-proportion Z-test, *p* = 0.460) nor OpenSpliceAI (*p* = 0.844) demonstrated a statistically significant improvement in accuracy over the original SpliceAI model. In contrast, the Legacy Ensemble performed significantly worse than all SpliceAI-based methods, with an accuracy of 78.5% (*p* < 0.001).

**Table 1:** Performance Metrics for CI-SpliceAI Benchmark.

|  | Sensitivity | Specificity | Balanced Accuracy |
| --- | --- | --- | --- |
| SpliceAI | 84.0% | 95.6% | 89.8% |
| CI-SpliceAI | 83.7% | 97.6% | 90.7% |
| OpenSpliceAI | 81.6% | 97.5% | 89.5% |
| Legacy Ensemble | 73.0% | 84.0% | 78.5% |

### Riepe Benchmark Results

Unlike the first analysis, this dataset revealed a more pronounced performance divergence among the models.

**Figure 2.**
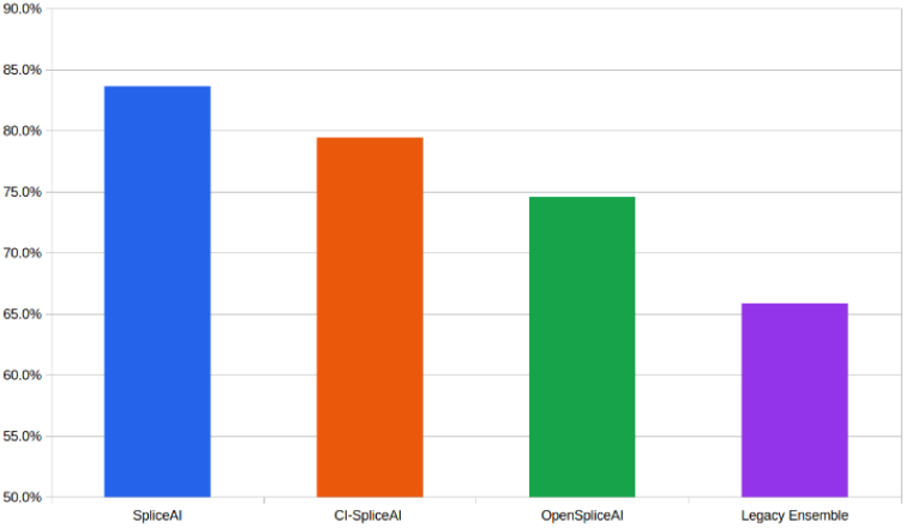
Balanced Accuracy on Riepe Benchmark

While the original SpliceAI achieved the highest balanced accuracy at 84.6%, CI-SpliceAI showed no statistically significant difference in accuracy (*p* = 0.219). However, OpenSpliceAI demonstrated a significantly lower balanced accuracy than SpliceAI at 74.6% (*p* = 0.019). All three algorithms significantly outperformed the Legacy Ensemble approach (*p* < 0.0001), which achieved a balanced accuracy of just 65.9%.

**Table 2:**
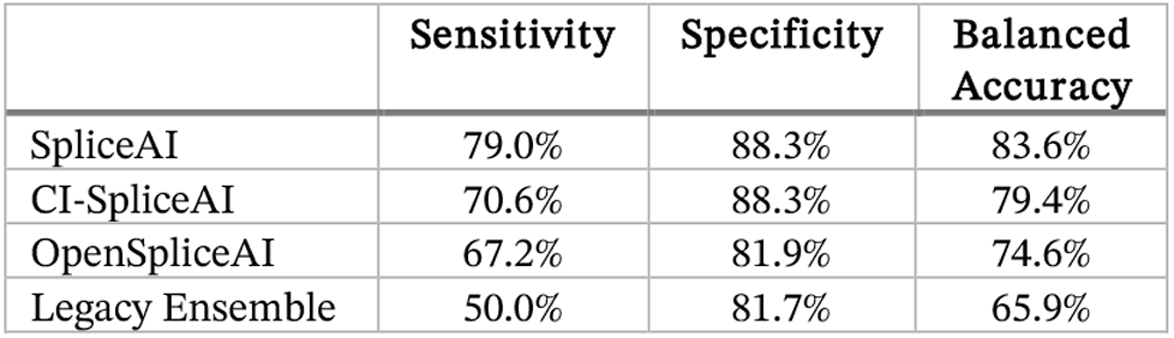
Performance Metrics for Riepe Benchmark.

|  | Sensitivity | Specificity | Balanced Accuracy |
| --- | --- | --- | --- |
| SpliceAI | 79.0% | 88.3% | 83.6% |
| CI-SpliceAI | 70.6% | 88.3% | 79.4% |
| OpenSpliceAI | 67.2% | 81.9% | 74.6% |
| Legacy Ensemble | 50.0% | 81.7% | 65.9% |

### ClinVar Benchmark Results

All three deep-learning algorithms demonstrated similarly high performance on the ClinVar Benchmark data, with only minor variations in accuracy.

**Figure 3.**
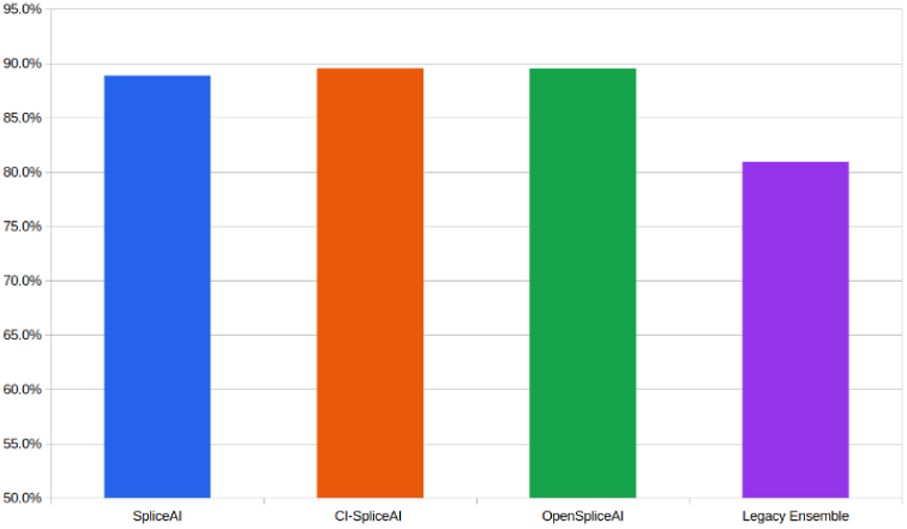
Balanced Accuracy on ClinVar Benchmark

Specifically, CI-SpliceAI and OpenSpliceAI achieved the highest balanced accuracy at 89.5%, demonstrating a small, but statistically significant, improvement when compared to the original SpliceAI algorithm (*p* < 0.001). Conversely, the Legacy Ensemble performed significantly worse than the other models, yielding the lowest balanced accuracy at 80.9% (*p* < 0.001).

**Table 3:** Performance Metrics for ClinVar Benchmark.

|  | Sensitivity | Specificity | Balanced Accuracy |
| --- | --- | --- | --- |
| SpliceAI | 79.1% | 98.7% | 88.9% |
| CI-SpliceAI | 80.0% | 99.1% | 89.5% |
| OpenSpliceAI | 79.9% | 99.2% | 89.5% |
| Legacy Ensemble | 83.2% | 78.7% | 80.9% |

### Correlation Analysis Results

#### Delta Score Correlation

Both open-source models demonstrated strong correlation with the original SpliceAI scores, as measured by Spearman’s rank correlation coefficient ρ. The degree of correlation varied by the type of splice event (Acceptor/Donor Loss/Gain), as shown in Table 4.

**Table 4:** Spearman’s Rank Correlation and Mean Absolute Difference on Delta Score.

|  | CI-SpliceAI |  | OpenSpliceAI |  |
| --- | --- | --- | --- | --- |
| Score | $\rho$ | MAD | $\rho$ | MAD |
| DS_AG | 0.786 | 0.070 | 0.668 | 0.077 |
| DS_AL | 0.864 | 0.029 | 0.924 | 0.024 |
| DS_DG | 0.808 | 0.069 | 0.677 | 0.079 |
| DS_DL | 0.883 | 0.025 | 0.940 | 0.022 |

CI-SpliceAI scores exhibited a strong correlation with original SpliceAI scores, ranging from ρ = 0.786 (DS_AG) to ρ = 0.883 (DS_DL). The mean absolute difference (MAD) was low across all categories, ranging from 0.025 to 0.07.

OpenSpliceAI also demonstrated strong correlations, with particularly high values for loss-of-function events (ρ = 0.924 for DS_AL and ρ = 0.94 for DS_DL). However, its correlation was noticeably lower for gain-of-function events (ρ = 0.668 for DS_AG and ρ = 0.677 for DS_DG) compared to CI-SpliceAI.

#### Delta Position Agreement

Analysis of the predicted splice site locations for variants with evidence of splicing disruption (delta score ≥ 0.2) showed strong agreement between the open-source models and the original SpliceAI algorithm, as indicated by the high exact match rate (EMR) across splice disruption types.

**Table 5:** Exact Match Rate for Delta Position.

|  | CI-SpliceAI | OpenSpliceAI |
| --- | --- | --- |
| Site Type | Exact Match Rate | Exact Match Rate |
| AG | 90.6%<br>(CI: 90.2 - 91.0%) | 92.5%<br>(CI: 92.1 - 92.9%) |
| AL | 96.1%<br>(CI: 95.7 - 96.4%) | 97.6%<br>(CI: 97.3 - 97.8%) |
| DG | 92.5%<br>(CI: 92.2 - 92.9%) | 95.5%<br>(CI: 95.2 - 95.8%) |
| DL | 97.5%<br>(CI: 97.2 - 97.8%) | 98.5%<br>(CI: 98.2 - 98.7%) |

Both models achieved an exact match rate of over 90% for all four splice categories, demonstrating that when a variant is predicted to disrupt splicing, the location of the affected site is nearly always consistent with the original SpliceAI algorithm. The exact match rate was consistently higher for OpenSpliceAI, ranging from 92.5% for DS_AG to its highest at 98.5% for DS_DL.

## Discussion

### Benchmark Analysis

These results indicate that both CI-SpliceAI and OpenSpliceAI successfully replicate the core predictive capabilities of SpliceAI, though with notable differences in performance characteristics across different evaluation contexts. On the CI-SpliceAI benchmark dataset, all three algorithms achieved balanced accuracies exceeding 89%, with no statistically significant differences in accuracy. Similarly, on the large-scale ClinVar benchmark comprising over 58,000 variants, CI-SpliceAI and OpenSpliceAI demonstrated nearly identical performance to the original algorithm, with balanced accuracies of 89.5% compared to SpliceAI’s 88.9%.

While performance was largely equivalent on the literature-derived and large-scale ClinVar datasets, the Riepe benchmark revealed more substantial differences between algorithms. On this dataset of functionally validated splice assays, the original SpliceAI achieved the highest balanced accuracy at 83.6%, followed by CI-SpliceAI at 79.4%, and OpenSpliceAI at 74.6%. The performance gap between SpliceAI and OpenSpliceAI reached statistical significance (p = 0.019), representing the only benchmark where one of the open-source implementations demonstrated meaningfully inferior performance. This divergence may be attributable to gene-specific splicing patterns in ABCA4 and MYBPC3 that are not fully captured by OpenSpliceAI’s training data.

### Score Correlation and Position Agreement

The correlation analysis provides additional nuance to our understanding of algorithm concordance. CI-SpliceAI demonstrated relatively balanced correlation across all splice event types, with Spearman’s ρ values ranging from 0.786 to 0.883. In contrast, OpenSpliceAI showed a striking asymmetry, with very high correlation for loss events (ρ = 0.924 for DS_AL and ρ = 0.940 for DS_DL) but substantially lower correlation for gain events (ρ = 0.668 for DS_AG and ρ = 0.677 for DS_DG).

This asymmetry suggests that OpenSpliceAI may have learned different feature representations for splice site gains compared to the original SpliceAI, despite achieving similar overall accuracy. The consistently lower mean absolute difference values for loss events across both algorithms (0.022-0.029) compared to gain events (0.069-0.079) indicates that predicting the disruption of existing splice sites is inherently more consistent across implementations than predicting the activation of cryptic splice sites.

Despite these differences in score distributions, both open-source implementations demonstrated remarkably high agreement with SpliceAI in predicting the locations of affected splice sites. Exact match rates exceeded 90% for all splice event types, reaching as high as 98.5% for donor loss events with OpenSpliceAI. This high spatial concordance is clinically significant, as accurately localizing the affected splice site is often as important as detecting its presence for understanding the disease mechanisms.

### Limitations of Sequence-Based Predictions

All three deep learning algorithms substantially outperformed the Legacy Ensemble approach, which combines four earlier-generation splice prediction tools. The Legacy Ensemble achieved balanced accuracies ranging from 65.9% to 80.9% across the three benchmarks, compared to 74.6% to 90.7% for the deep learning methods. This performance gap demonstrates the transformative impact of deep learning on computational splice prediction.

However, none of the algorithms in our study achieved a balanced accuracy greater than 90.7%, leaving substantial room for improvement. This performance ceiling reflects a fundamental limitation of sequence-based prediction approaches. Because alternative splicing is influenced not only by the underlying RNA sequence but also by the multiple regulatory interactions of various molecular factors within the cell [12], any splice site prediction algorithm based strictly on the DNA sequence will face an asymptotic limit to its maximum achievable accuracy.

### Considerations for Algorithm Selection

Based on these findings, CI-SpliceAI represents the optimal choice among open-source alternatives. It demonstrated consistent performance across all three benchmarks without statistically significant differences from the original SpliceAI, while exhibiting more balanced correlation patterns across different splice event types compared to OpenSpliceAI. The use of collapsed isoforms from GENCODE in training also provides theoretical advantages for detecting alternative splicing events, even if these advantages were not definitively demonstrated in our benchmarks.

OpenSpliceAI remains a viable alternative, particularly for applications that prioritize the detection of splice site losses, where it demonstrates the strongest correlation with the original algorithm. However, its relatively weaker performance on the Riepe functional validation dataset and lower correlation for gain events suggest some caution is warranted for applications requiring the highest possible accuracy.

For organizations already using the original SpliceAI with access to its precomputed score databases, there is little motivation to adopt an alternative implementation on the basis of accuracy alone. However, commercial clinical laboratories face several practical considerations that extend beyond predictive performance when selecting a splice prediction solution.

First, licensing constraints are a critical barrier. The original SpliceAI model weights and precomputed scores are restricted to academic, non-commercial use, preventing their integration into validated clinical pipelines. This alone necessitates alternative implementations for commercial laboratories.

Second, computational throughput must be considered. Given the volume of variants processed in clinical settings, it is generally infeasible to run deep-learning models on every variant in real time. Instead, laboratories must utilize precomputed genome-wide scores for efficient prediction. At present, no official precomputed scores exist for either CI-SpliceAI or OpenSpliceAI, limiting their immediate clinical utility despite the advantage provided by their permissive licensing.

Finally, there are significant limitations associated with the existing SpliceAI precomputed scores. The widely used Illumina-provided datasets were generated only for the GRCh37 reference genome, and the available GRCh38 scores were derived through an error-prone liftover process rather than native computation. In addition, these precomputed scores rely on outdated transcript annotations that no longer reflect current gene models, particularly for genes with extensive alternative splicing.

To address these gaps, new genome-wide splice prediction scores for these alternative methods must be computed natively for both GRCh37 and GRCh38 using up-to-date transcript definitions. Such resources are essential for enabling the adoption of these open-source solutions in high-throughput clinical environments.

### Study Limitations

Several limitations of this study should be acknowledged. First, the Riepe benchmark dataset, while consisting of gold-standard functionally validated variants, includes only 213 variants from two genes. Larger functional validation datasets would strengthen confidence in the observed performance differences. Second, the ClinVar benchmark dataset, while covering a large spectrum of variation, relies on clinical classifications rather than direct functional validation and includes variants with varying levels of evidence quality. Finally, our study focused exclusively on splice site prediction and did not evaluate other aspects of algorithm performance, such as computational efficiency, ease of use, or applicability to non-human species.

## Conclusion

In this study, we evaluated the performance of three different deep-learning algorithms for splice prediction. Our results indicate that the two open-source reimplementations of SpliceAI successfully replicate the predictive performance of the original algorithm across multiple evaluation contexts. CI-SpliceAI and OpenSpliceAI both achieve high balanced accuracy for splice site prediction and represent viable alternatives to the original SpliceAI algorithm, particularly for applications requiring open-source licensing or custom model training. The availability of these high-quality open-source tools democratizes access to state-of-the-art splice prediction and facilitates continued innovation in this important area of genomic medicine.

## Supporting information

ClinVar Benchmark Dataset

## Notes

### Competing Interest Statement

The authors have declared no competing interest.

